# Reducing Cholesterol in Macrophage Activates NF-kB through Mitochondria, Resulting in Epigenomic Reprogramming to Dampen Inflammation

**DOI:** 10.1101/2022.02.10.479926

**Authors:** Zeina Salloum, Kristin Dauner, Kiran Nakka, Neha Verma, David Valdivieso-González, Víctor Almendro-Vedia, Jeffery McDonald, Hina Bandukwala, Alexander Sorisky, Iván López-Montero, Jeffery Dilworth, Xiaohui Zha

## Abstract

Cholesterol plays an important role in macrophage functions including their immune response^1^. Recently, NF-*k*B was shown to reprogram the epigenome in macrophages^2^. Here, we show that NF-kB pathway is activated in resting macrophages when cholesterol is reduced by statin or methyl-β-cyclodextrin (MCD). Activated NF-*k*B increases the expression of histone-modifying enzymes, such as demethylase JMJD3. We provide evidence that the epigenome in these macrophages is reprogrammed, likely driven by NF-*k*B and histone modifications^2^. We also show that cholesterol reduction in macrophages results in suppression of mitochondria respiration. Specifically, cholesterol levels in the inner membrane of the mitochondria is reduced, which impairs the efficiency of ATP synthase (complex V). Consequently, protons accumulate in the intermembrane space to active NF-*k*B and JMJD3, thereby modifying the epigenome. When subsequently challenged by the inflammatory stimulus lipopolysaccharide (LPS), cholesterol-reduced macrophages generate responses that are less pro-inflammatory and more homeostatic, which should favour inflammation resolution. Taken together, we describe a mechanism by which the level of mitochondrial cholesterol in resting macrophages regulates the epigenome through NF-kB, thereby preparing macrophage for future immune activation.

---

Macrophages primarily function to defend against pathogens and danger signals by quickly activating a large transcriptome. This transcriptome is mainly defined by lineage-determining transcription factors (TFs), but also relies on signal-dependent TFs to trigger large-scale gene expression from an epigenetically configurated genome. Such epigenome configurations reflect metabolic and environmental cues, including those from past events, thereby fine-tuning context-specific inflammatory responses in macrophages^3^.

Many chronic diseases, such as atherosclerosis, are associated with a low-grade level of inflammation^4^. In atherosclerosis, macrophages in the lesions are overloaded with cholesterol and become inflamed^5^. It is not clear how macrophages respond to environmental factors, for instance cholesterol content, to modulate their transcriptome that in turn determines their specific inflammatory responses. Previously, we and others have observed that the level of cholesterol in resting macrophages directly correlates with the pattern of inflammatory activation^6-8^. For example, when encountering LPS, cholesterol-overloaded macrophages release more inflammatory cytokines, such as TNF-α, IL-6, and IL12p40, and fewer anti-inflammatory cytokines such as IL-10, in comparison to control macrophages. Conversely, macrophages with less cholesterol express fewer inflammatory cytokines and more IL-10 upon identical LPS exposure, compared to control macrophages^6^. LPS activates the expression of several hundred genes within a short period of time^9^. This is made possible by using relatively few signal-dependent TFs on open (or partially open) genome regions, which is the result both lineage-determining TFs^10^ and metabolic/environmental factors^11^. Cholesterol is known to influence LPS responses in macrophages^12^, potentially by modulating the genome into a poised epigenomic state, thus fine-tuning gene expression in inflammatory responses^13, 14^.

To understand if cholesterol levels alone can influence genomic function in resting macrophages, we treated macrophages with MCD for 1 h or statin for 2 days, which reduces cellular cholesterol by about 30%^6, 15^ and performed RNA-seq at the end of treatments. We found that 264 and 2483 genes are upregulated in macrophages treated with MCD or statin, respectively, with 218 overlapping genes (Fig. 1 A). To study the potential mechanisms, we analyzed major TFs responsible for the cholesterol-lowering induced gene expression, using a database of mouse transcriptional regulatory networks^16^ embedded in Metascape^17^. Analysis of promoter sequences at differentially expressed genes identified Nfkb1 as the top regulator for genes upregulated by both MCD and statin (Fig. 1 B, *a & b*). Nfkb1 is a major signal-dependent TF for gene expression during inflammatory responses in activated macrophages^18^. We also performed RNA-seq in LPS-activated macrophages and similarly analyzed TF regulatory networks. LPS triggers the expression of 2343 genes within 3 h. The analysis of TF regulatory pathway identified Nfkb1 as the top regulator, along with other major inflammatory regulatory TFs, such as Stat1 & 3, IRFs, JUN et al (Fig. 1 C, *a*). Comparing genes induced by MCD or statin in resting macrophages with those induced by LPS, we found that 15 out of the 20 top regulators of LPS activated genes were shared between the treatments (Fig. 1 C, *b*). This similarity is surprising but, however, may imply some overlapping mechanisms. Nevertheless, the upregulation of NF-*k*B target genes (such as IL-1β and TNF-α) by MCD or statin is relatively modest in comparison with their activation by LPS (Fig. 1 D, *a & b*), indicating a substantial difference in the magnitude of NF-*k*B activation. To confirm the role of NF-*k*B in mediating the effects of cholesterol depletion, we observed that blocking NF-*k*B function, using two structurally unrelated inhibitors, prevented the upregulation of IL-1β and TNF-α expression in MCD-treated macrophages (Fig. 1 E, *a & b*). We thus conclude that cholesterol reduction in resting macrophages by MCD activates NF-*k*B pathways, similar but to a lesser extent than LPS-stimulated macrophages.

**Figure 1.**
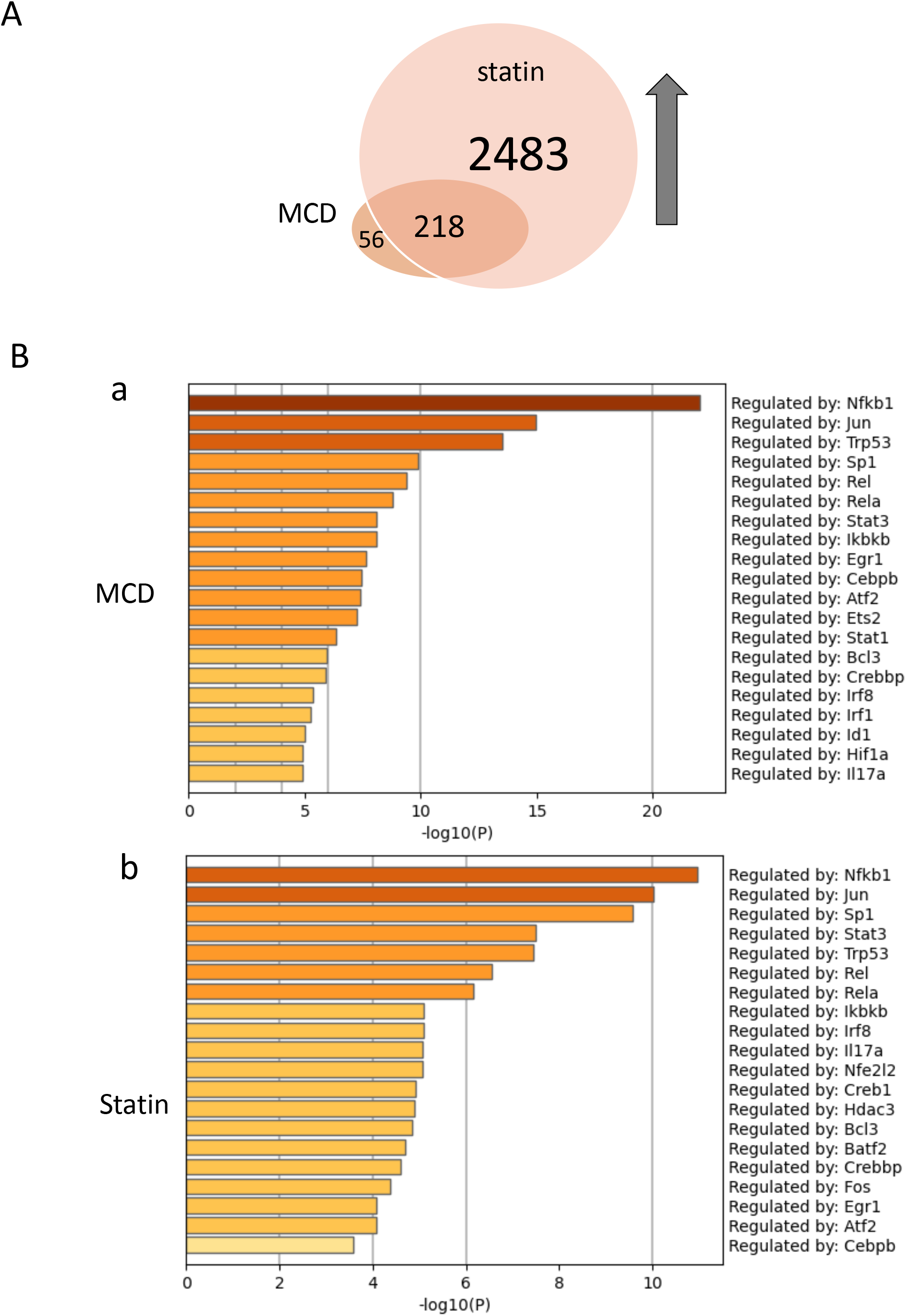
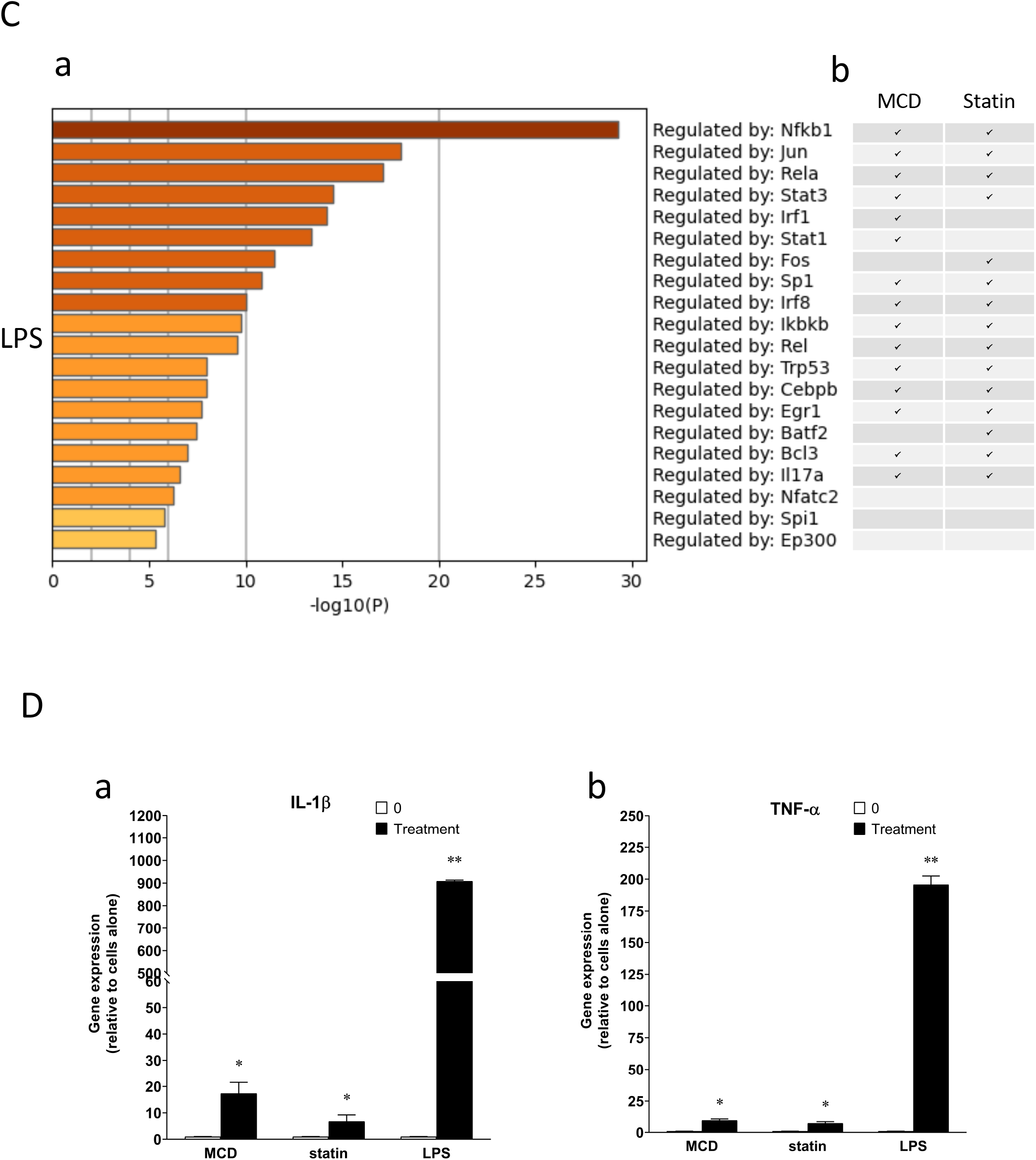
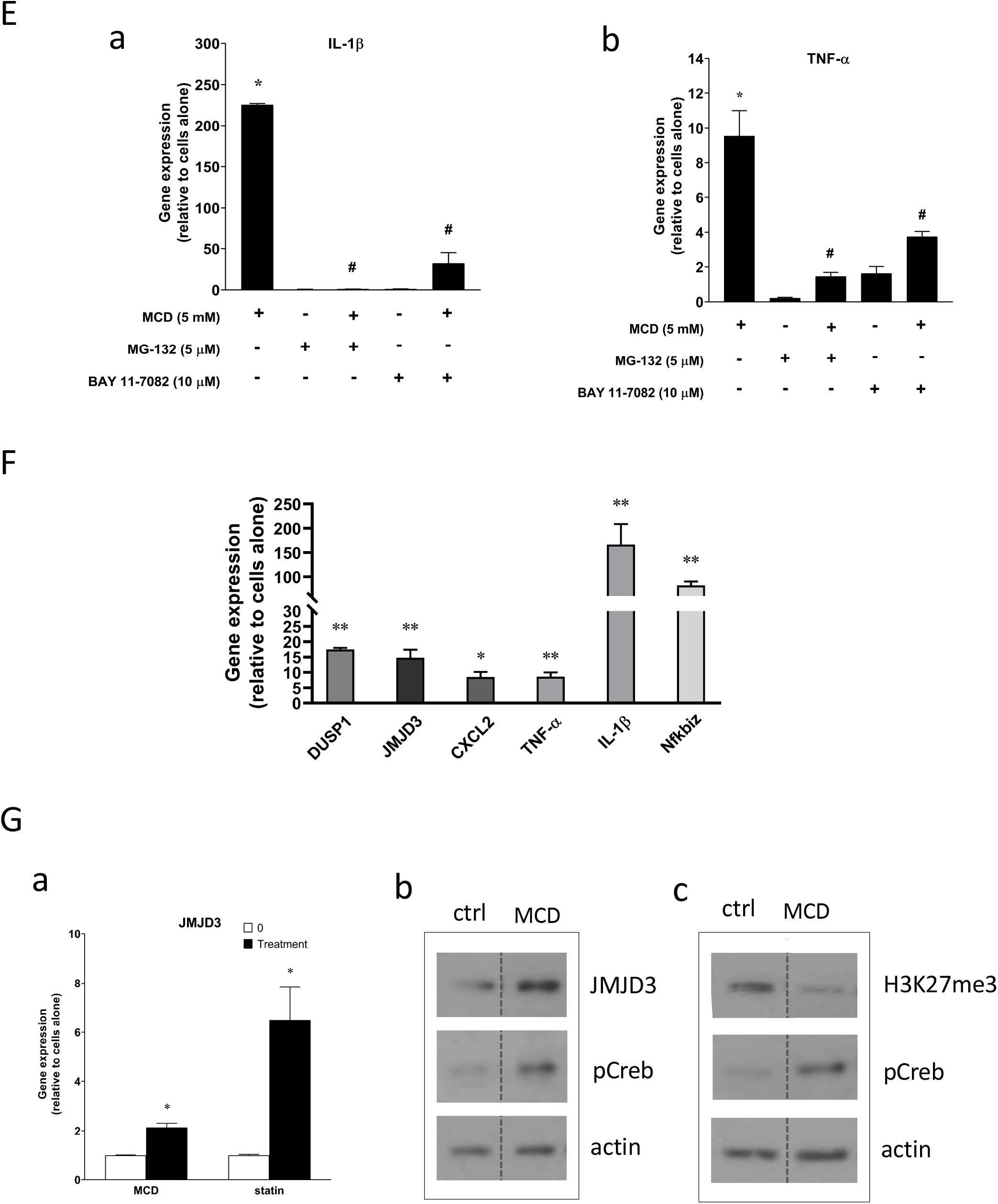
Cellular cholesterol contents regulate NF-kB pathway. (**A**) Venn diagram on RNA-seq datasets of macrophages with or without MCD (5 mM; 1-h), statins (7 μM; 2 days). (**B**) TF regulators (TFs) of activated genes were analyzed using Metascape program^37^, by either MCD (**a**) or statins (**b**); (**C**) TFs LPS-stimulated genes LPS (100 ng/ml; 3 hours); (**a**) Top 20 TFs upregulated by LPS; (**b**) overlaps of LPS-upregulated TF regulatory pathways with that by MCD or statins. (**D**) RT-qPCR of IL-1β (**a**) and TNF-α (**b**) in macrophages with or without MCD. (**E**) Effect of NF-kB inhibitors (MG-132 [5 μM] and BAY11-7082 [10 μM]) on IL-1β (**a**) and TNF-α (**b**) expression in MCD-treated cells. (**F**) MCD upregulates NF-kB target genes. (**G**) JMJD3 gene expression in MCD, or statins treated macrophages (**a**); western blot of JMJD3 protein (**b**) and H3K27Me3 (**c)** in macrophages treated with 5 mM MCD (1h). pCREB was used as internal control for cholesterol depletion^6^ and actin a loading control. Bars are representative of 3 independent experiments with 3 replicates and are presented as means ± SD. Statistical analysis was performed using unpaired, two-tailed Student’s t-test. An asterisk (*) or double asterisks (**) indicate a significant difference with p<0.05 and p<0.001, respectively. A hashtag (#) indicates a significant difference between MCD-treated under a specific condition (drug effect) with p < 0.05.

We verified several NF-*k*B target genes in MCD-treated macrophages using qRT-PCR (Fig. 1, F). Among these, expression of the histone H3K27-demethylase JMJD3 is upregulated by both MCD and statin treatments (Fig. 1 G, *a*). This is consistent with previous results showing that JMJD3 is a NF-kB target gene^19^. We confirmed that JMJD3 protein level is increased (Fig. 1 G, *b*) and the global levels of H3K27me3, the substrate of JMJD3, are decreased by MCD (Fig. 1 G, *c*). Similar results were seen in macrophages treated with statin (Supplementary Fig. 1 A). In addition to JMJD3, H3K27me3 can also be demethylated by UTX, encoded by KDM6a. However, KDM6a is not upregulated by MCD (Supplementary Fig. 1 B). A reduction in H3K27me3 could also be due to a decrease in one of the H3K27 methyltransferases, Ezh1 or Ezh2, but neither is downregulated by MCD (Supplementary Fig. 1 C). This suggests that MCD/statin treatment activates NF-kB, which in turn induces the expression of JMJD3 to demethylate H3K27me3. Again, the expression of JMJD3 is upregulated significantly more in LPS-treated macrophages than those treated by MCD (Supplementary Fig. 1 D), in agreement with the difference in magnitude of NF-kB activation on IL-1β and TNF-α expression shown in Fig. 1 D.

Recently, NF-kB activation was shown to reprogram the epigenome through nucleosome displacement, which increases the accessibility NF-*k*B-target genes for transcription^2^. NF-*k*B is also shown to trigger histone modification to further alter/stablize the epigenome. In fact, it is histone modification that ensures the epigenome remains reprogrammed long after initial NF-kB activation^2^. We therefore investigated the effect of cholesterol reduction on the macrophage epigenome. Resting macrophages, treated with or without MCD (1 h), were analyzed for the assay for transposase-accessible chromatin with sequencing (ATAC-seq). As shown in Fig. 2 A, lowering cellular cholesterol by MCD significantly altered the epigenome: 2791 genes more open (red), whereas ~ 1000 genes were less open (grey). Focusing on genomic regions that became more open after MCD treatment, we performed TF regulator network analysis^17^. As in RNA-Seq, we observed that Nfkb1 was the top ranked TF regulator for genomic loci that were opened upon MCD treatment. In fact, much of the enriched TF ranking resembled the those from LPS RNA-seq analysis, i.e. mainly inflammation-related (Fig. 2 B). Noticeably, Nfkb1- and Rel *a*-regulated genes occupy predominant spots on the volcano plot (Fig. 2 A). Aligning the RNA-seq and ATAC-seq tracks for NF-kB target genes IL-1β and TNF-α showed the two genes are more open in MCD treated macrophages, particularly at promoter regions (Fig. 2 C, ATAC-seq). This opening of chromatin at these two genes coincides with increased expression in MCD- or statin-treated macrophages (Fig. 2 C, RNA-seq).

**Figure 2.**
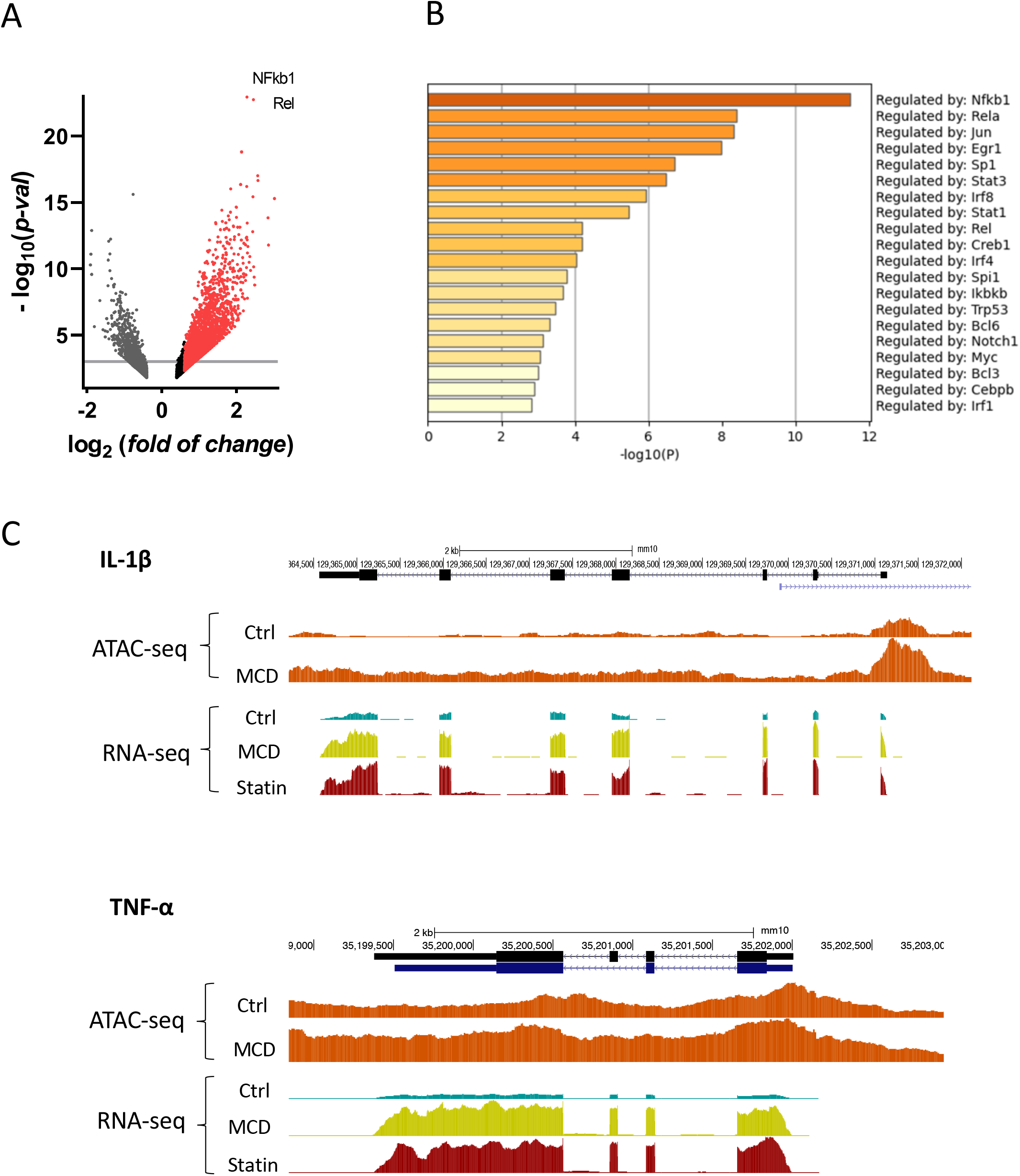
Cellular cholesterol contents regulate epigenetic landscape. (**A**) Volcano plot of differentially accessible peaks from ATAC-seq between MCD-treated and untreated macrophages (**B**) TF regulator pathway analysis of ATAC-seq for more opened genes (red)^39^; (**C**) ATAC-seq and RNA-seq profiles alignment from ATAC-seq and RNA-seq for the genomic loci of IL-1β (top) and TNF-α (bottom).

This similarity of the expression profile in RNA-seq in MCD- or statin-treated resting macrophages with that of LPS-activated macrophages (Fig. 1) was unexpected. In recent years, it has become evident that, when LPS triggers inflammatory responses, there is a dramatic metabolic shift in the mitochondria from the oxidative phosphorylation (OXPHOS) towards glycolysis^20^. This shift is essential for macrophage inflammatory activation^21^. In fact, the LPS/TLR4 complex was observed to physically interact with the mitochondria and modulate mitochondria organization^22^. We therefore investigated if lowering cholesterol also influences mitochondria OXPHOS in resting macrophages. Using Seahorse extracellular flux analyzer, we determined the oxygen consumption rates (OCR) of macrophages treated with or without MCD (Fig. 3 A, *a*). We found that MCD treatment decreases overall resting mitochondrial OCR (Fig. 3 A, *b*), which is almost entirely from oligomycin-sensitive component, the ATP synthase, a component of the electron transport chain (ETC) in the inner mitochondrial membrane (IMM) (Fig. 3 A, *c*). Importantly, the maximal respiration is unchanged (Fig. 3 A, *d*). Therefore, MCD treatment of resting macrophages specifically suppresses mitochondria ATP synthase. Statin treatment also decreased mitochondrial OCR parameters and broadly supresses all ETC, including ATP synthase (Supplementary Fig. 2 A & C). Together, we conclude that cholesterol levels in macrophages influence mitochondria respiration.

**Figure 3.**
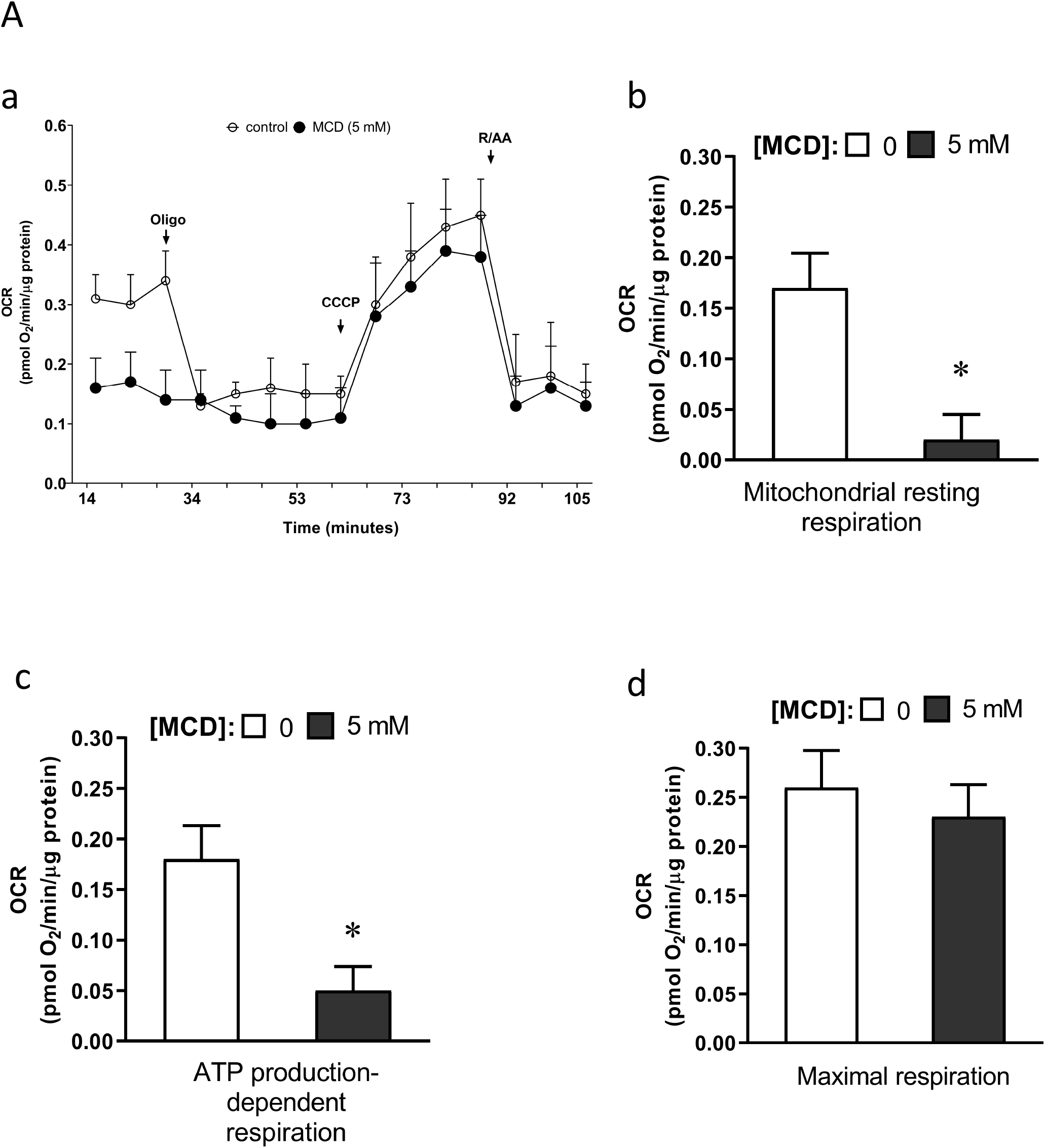
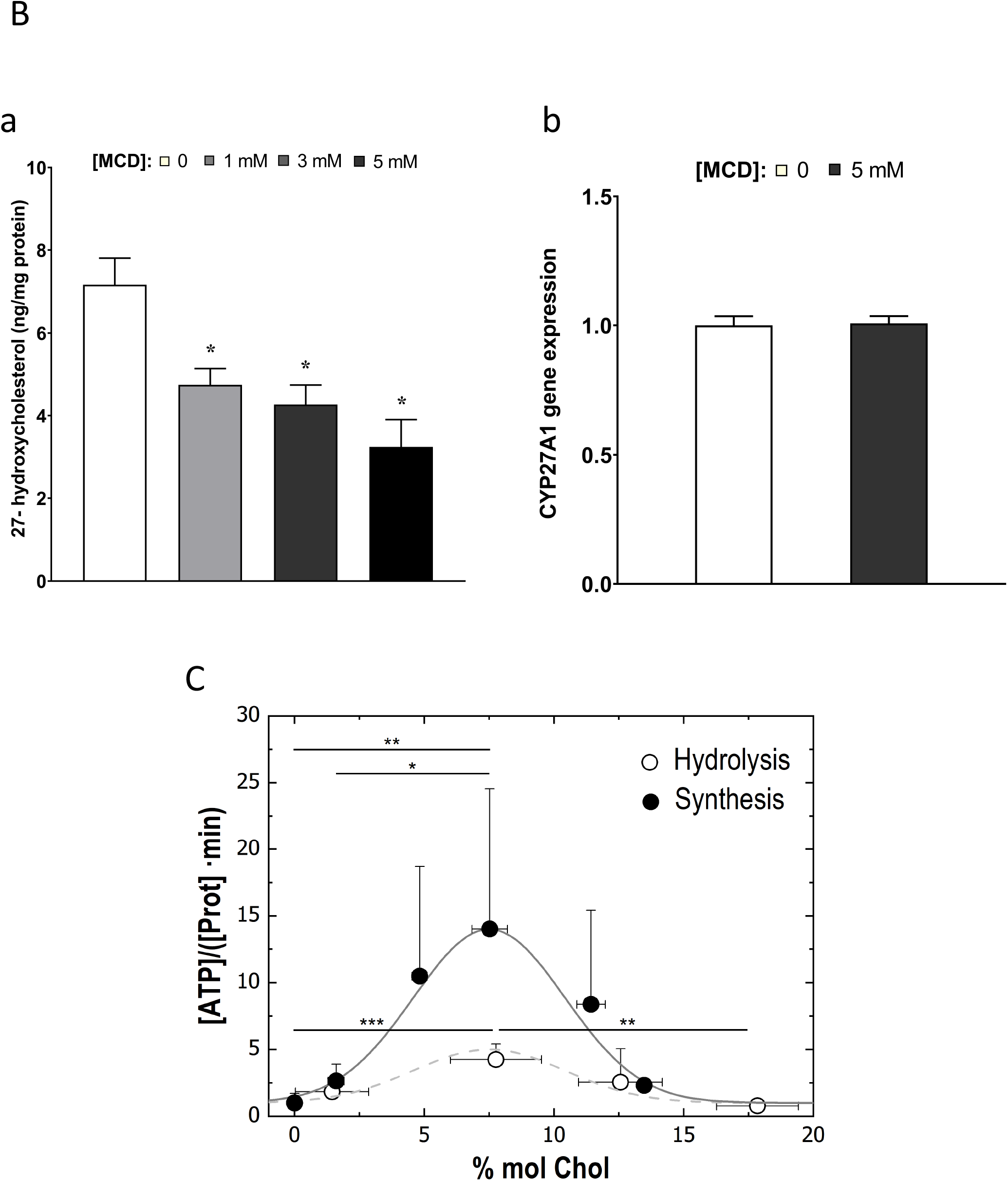
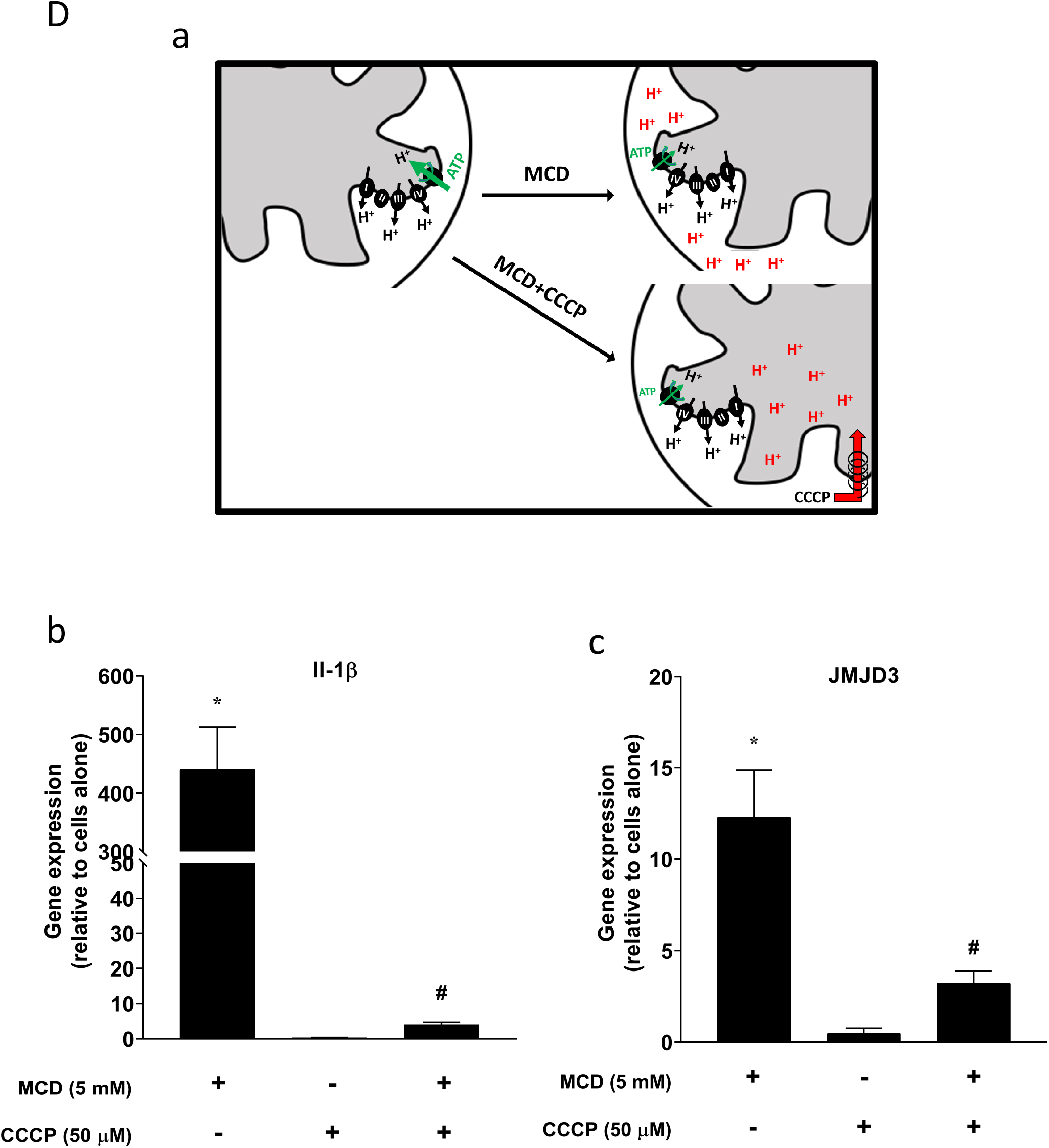
Cholesterol modulates mitochondria to activate NF-kB pathway. (**A**) (**a**) Mitochondrial oxygen consumption rates (OCR); (**b**) OCR for mitochondrial resting respiration; (**c**) OCR representing mitochondrial ATP production-linked respiration; (**d**) OCR representing maximal respiration of macrophages (+/- 5 mM MCD). (**B**) 27-hydroxycholesterol (27-HC) production: (**a**) 27-HC analysis by ultraperformance liquid chromatography/electrospray ionization/tandem mass spectrometry. 27-HC levels were normalized to the protein content in the whole cell pellet; (**b**) sterol 27-hydroxylase, CYP27A1, gene expression in MCD-treated cells and control cells. (**C**) ATP hydrolysis and synthesis in cholesterol-doped inner membrane vesicles (IMVs); ATP hydrolysis was performed by adding a total concentration of 2 mM ATP to 200 mM IMVs (lipid concentration) and incubated for 30 minutes. Concentration of phosphates from ATP hydrolysis were measured using the malaquite green assay^39^. ATP concentration after synthesis was measured using ATP detection assay kit (Molecular Probes) with a luminometer GloMax®-Multi Detection. (**D**) Effect of proton flux on NF-kB activation in MCD-treated cells. Schematic of potential mechanism induced by MCD, and MCD/CCCP involving mitochondrial proton flux (**a**); effect of CCCP (50 μM) on Il-1β (**b**) and JMJD3 (**c**) expression in MCD-treated cells. Data in (A) are representative of three independent experiments and presented as means± SD. Data in (B-E) are representative of 3 independent experiments with 3 samples per group and data are presented as mean ± SD. Statistical analysis was performed using unpaired, two-tailed Student’s t-test. An asterisk (*) and a hashtag (#) indicate a significant difference with p < 0.05.

We considered how MCD might influence mitochondria function, particularly through a specific component of the electron transport chain (ETC), i.e. ATP synthase in the IMM. In mammalian cells, the majority cholesterol is in the plasma membrane^23^. However, all intracellular membranes, including those in the mitochondria, also contain cholesterol^23, 24^. At molecular level, MCD chelates cholesterol from the plasma membrane without directly interacting with, or entering the cells^25^. However, cholesterol is highly exchangeable among cellular membranes and, as such, its distribution among membranes is governed by a dynamic steady state^23^. A reduction in cholesterol in the plasma membrane created by MCD would cause intracellular cholesterol to flow out to the plasma membrane to re-establish a new steady state. MCD therefore rapidly decreases cholesterol not just in plasma membrane, but also in intracellular membranes, including those in mitochondria^24^. Levels of IMM cholesterol can be determined by the activity of sterol 27-hydroxylase, an IMM resident enzyme. It catalyzes the conversion of cholesterol to 27-hydroxycholesterol and the rate is governed by substrate (cholesterol) availability in IMM^26^. Therefore, the amounts of 27-hydroxy cholesterol in macrophages directly reflect cholesterol levels in IMM^24^. Using mass spectrometry, we found that cellular 27-hydroxycholesterol content is decreased in a dose-dependently manner after a 1 h MCD treatment (Fig. 3 B, *a*). The expression level of sterol 27-hydroxylase, encoded by CYP27A1, is unchanged by 1 h MCD treatment (Fig. 3 B, *b*). We conclude that, by removing cholesterol from the plasma membrane, MCD effectively lowers cholesterol levels in IMM^*^. This could affect the activities of ETC components embedded in IMM, particularly complex V, ATP synthase.

To understand precisely how cholesterol contents in IMM may influence ATP synthase activities, we turned to an *in-vitro* membrane system composed of inner membrane vesicles from *Escherichia coli*^27^ (IMVs). *E. coli* inner membrane shares common features with IMM in mammalian cells^28^, including ATP synthase functions^29^; but lack cholesterol or any other sterol derivative in their lipid composition^30^. Various levels of cholesterol were then incorporated into the IMVs^31^. We observed that the activity of the complex V, both in the ATP synthesis and hydrolysis mode, is highly sensitive to cholesterol levels in the membrane: the highest activity is found in membranes with about 7% cholesterol; lower cholesterol levels in the membranes decrease ATP synthase activities (Fig. 3 C). These observations are consistent with known steady-state IMM cholesterol contents (about 5% of phospholipids)^32^. The *in-vitro* experiment therefore supports the notion that MCD lowers cholesterol in the IMM (Fig. 3 B, *a*), which in turn suppresses ATP synthase activity (Fig. 3 A, *c*).

ATP synthase in IMM utilizes proton flow from the inner space to the matrix to generate ATP. As shown by extracellular flux analysis (Fig. 3 A), MCD suppresses the activities of ATP synthase (thus fewer protons should flow down into the matrix), whereas the rest of ETC is largely unchanged (normal proton pumping into the inner space). Consequently, more protons would accumulate in the inner space in MCD-treated macrophages (Fig. 3 D, a), which in turn could result in the activation of NF-kB. Indeed, LPS is known to suppress mitochondrial ATP synthesis and thus to increase the proton gradient to active NF-kB^33^. To test this possibility, we treated macrophages with MCD in the presence of carbonyl cyanide m-chlorophenyl hydrazine (CCCP), a mitochondria proton ionophore that prevents proton buildup in the inner space^34^. In the presence of CCCP, MCD fails to upregulate IL-1β or JMJD3 (Fig. 3 D, *b & c*). We also tested another structurally unrelated mitochondrion proton ionophore BAM15^35^. It also abolished MCD-induced IL-1β or JMJD3 expression (supplementary Fig. 3, *a & b*). Taken together, we conclude that, by removing cholesterol from the plasma membrane, MCD effectively decreases cholesterol levels in IMM. This reduced cholesterol in IMM suppresses complex V activity, leading to proton buildup in the inner space of the mitochondria, which induces NF-kB activation, thereby resulting in increased IL-1β/JMJD3 expression.

Activated NF-kB was recently shown to displace nucleosomes and thereby rendering them permissive to transcription^2^. NF-kB also changes the histone methylation status to maintain reprogrammed epigenome even after a stimulus is removed^2^. As shown above, cholesterol reduction in resting macrophages activates NF-kB, indicated by IL-1β and TNF-α expression, and modifies histone methylation through JMJD3 to create a cholesterol-specific epigenome. Such an epigenome could poise the macrophage epigenome for cholesterol-specific immune responses. To test this, we challenged MCD-treated macrophages with LPS. We found that pre-treatment of macrophages with MCD for 1 h, prior to LPS stimulation, suppressed the expression of about 400 LPS activated genes, which predominantly represents genes involved in inflammatory pathways^19^ (Fig. 4 A, *a*). At the same time, MCD attenuated the suppression in another 400 of the genes suppressed by LPS. Genes that show this more attenuated suppression are mostly involved in homeostatic cell functions (Fig. 4 A, *b*). This is consistent with the notion that LPS not only activates inflammatory factors, but also suppresses certain homeostatic cell functions, such as proliferation^21^. The overall effect of prior MCD treatment on LPS responses can be observed by heatmaps (Fig. 4 A, *c*). LPS triggers expression of a large group of genes in untreated macrophages (Fig. 4 A, *c*, **i & iii**; first two columns; LPS vs. control). Macrophages pre-treated with MCD also respond to LPS to express these genes, but in a lower magnitude (Fig. 4 A, *c*, **i**; third column). This gene group is mostly pro-inflammatory as shown in Fig. 4 A, *a*. On the other hand, for genes normally suppressed by LPS, pre-treatment by MCD weakens the suppression (Fig. 4 A, *c*, **ii**). For a small group of LPS stimulated genes (<200), macrophages pre-treated with MCD express higher levels compared to LPS treatment alone (Fig. 4 A, *c*, **iii**). This enhancement by MCD on LPS-stimulated genes indicates that MCD does not act by suppressing overall LPS/TLR4 signaling.

**Figure 4.**
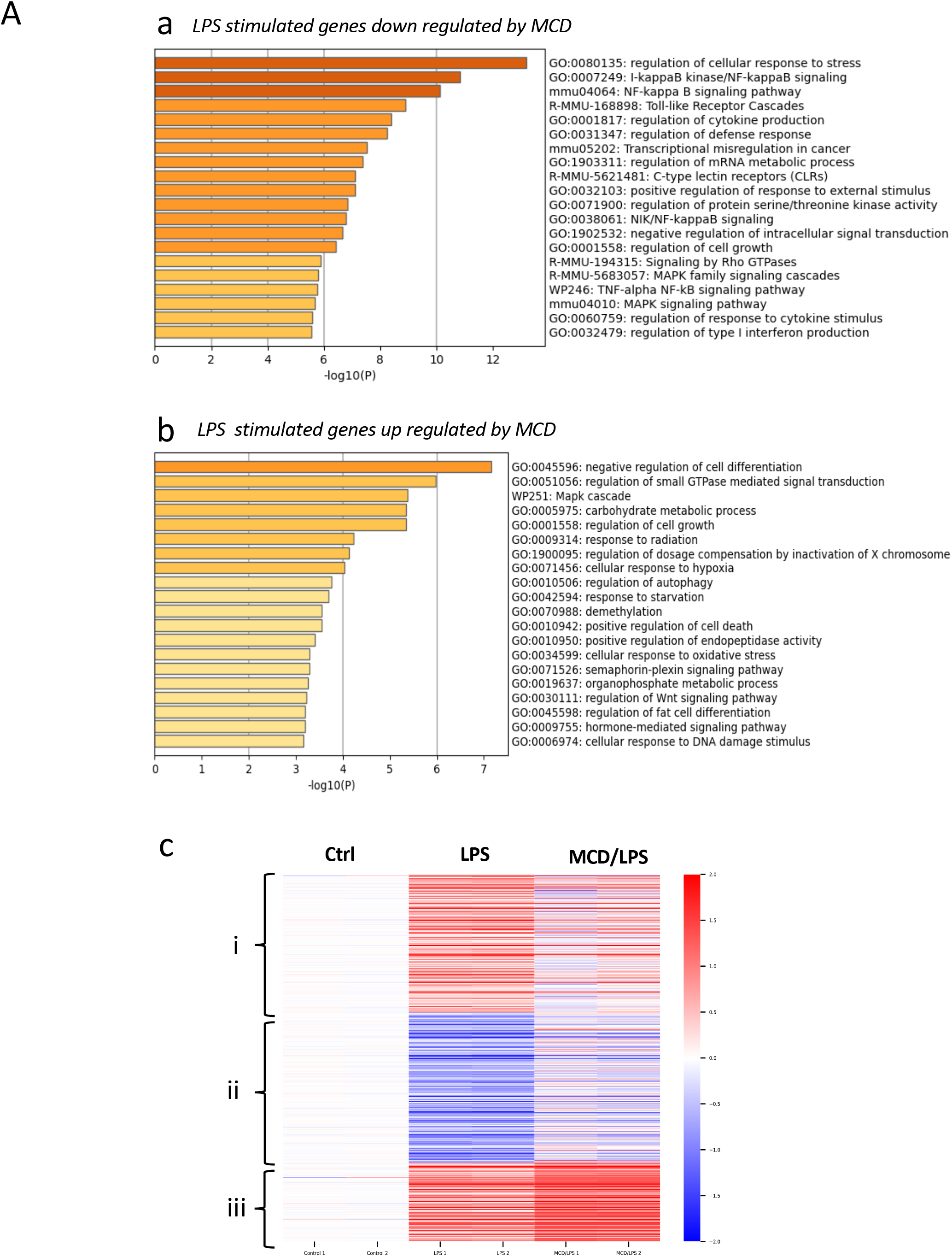
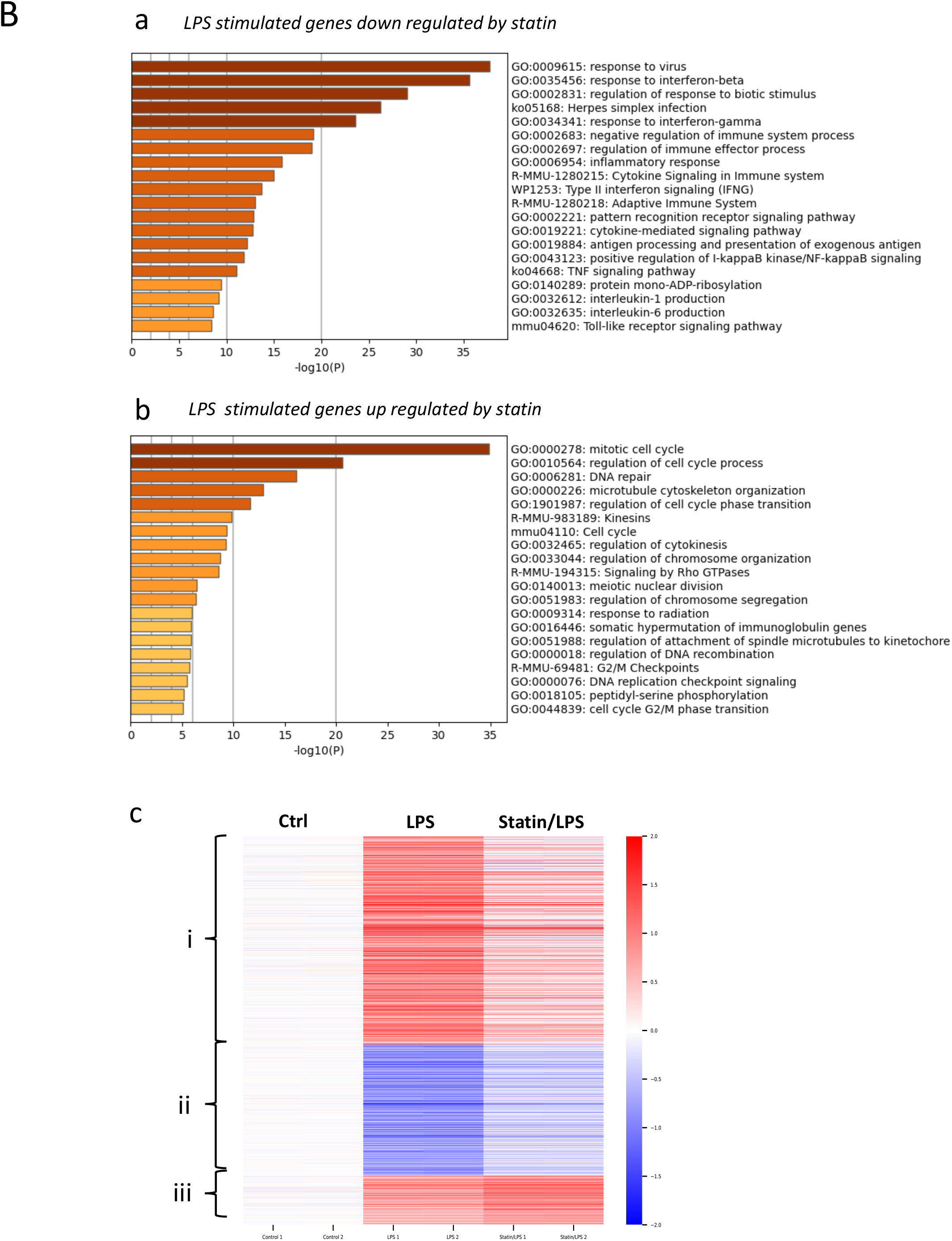
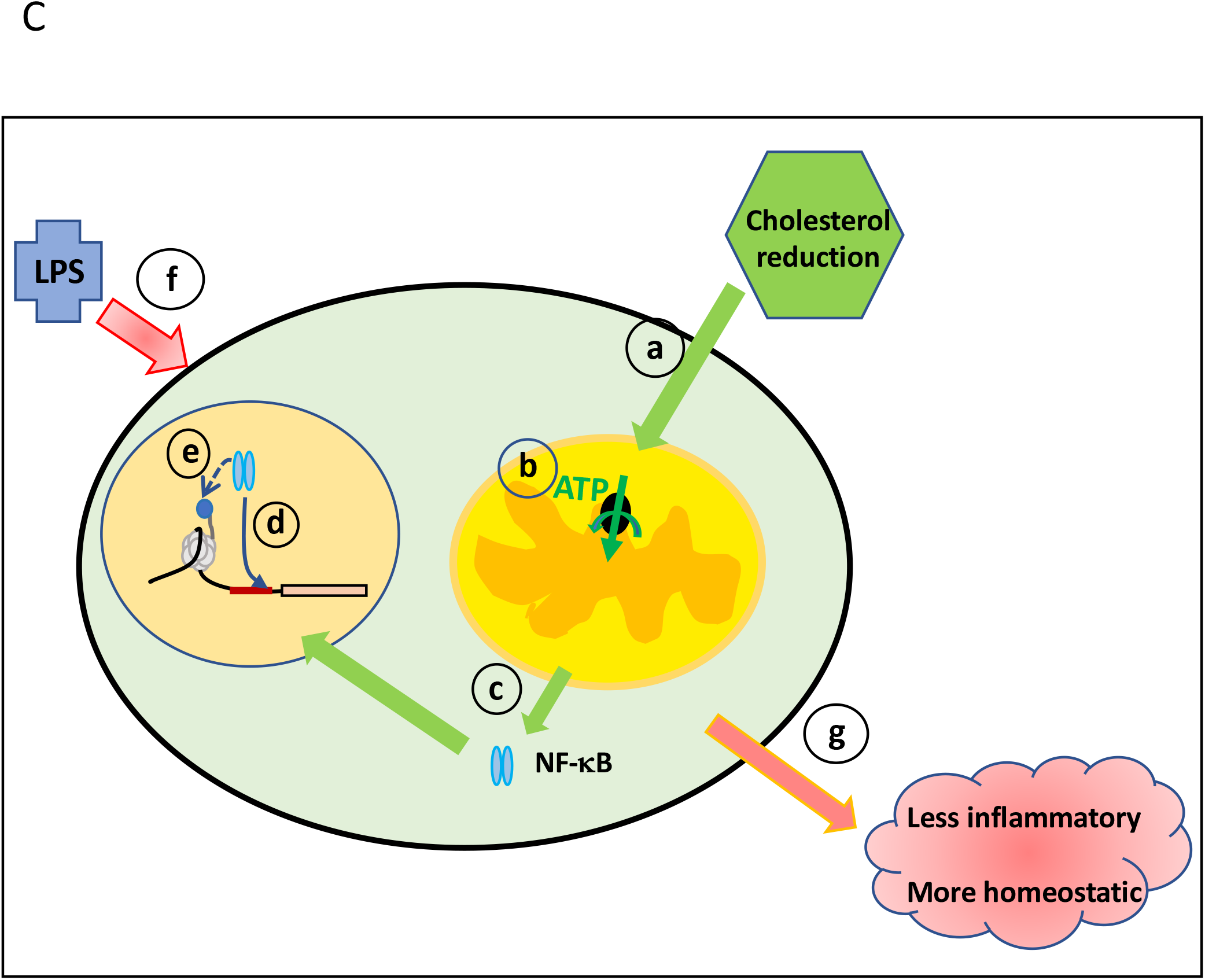
Cholesterol modulates mitochondria to reprogram LPS response. **(A)** Macrophages treated with MCD (1h) or without were stimulated with LPS (100ng/ml) for 3 h. (**a**) GO analysis of LPS-stimulated genes that downregulated by MCD; (**b**) GO analysis of LPS-stimulated genes that upregulated by MCD. (**c**) Heatmaps of gene expression in control, LPS and MCD/LPS treated macrophages: (**i**) genes upregulated by LPS, but less upregulated by statin/LPS; (**ii**) genes down regulated by LPS, but less down regulated by statin/LPS; (**iii)** genes upregulated by LPS and more upregulated by statin/LPS. **(B)** Macrophages treated with statin (2 d) or without were stimulated with LPS (100ng/ml) for 3 h. (**a**) GO analysis of LPS-stimulated genes that down regulated by statin; (**b**) GO analysis of LPS-stimulated genes that upregulated by statin. (**c**) Heatmaps of gene expression in control, LPS and statin/LPS treated macrophages: (**i**) genes upregulated by LPS, but less upregulated by statin/LPS; (**ii**) genes down regulated by LPS, but less down regulated by statin/LPS; (**iii)** genes upregulated by LPS and more upregulated by statin/LPS. (**C**) model of cholesterol modulating mitochondria to reprogram LPS responses.

An identical pattern is seen in macrophages treated with statin prior to LPS stimulation. Statin suppresses the expression 1023 LPS-activated genes in inflammatory pathways^19^ (Fig. 4 B, *a*) and, for genes suppressed by LPS, statin attenuates the suppression of more than 600 genes in homeostatic cell functions (Fig. 4 Fig. 4 B, *b*). Again, heatmaps (Fig. 4 B, *c*) show that statin suppresses LPS-stimulated pro-inflammatory genes (Fig. 4 B, a, and 4 B, *c*, **i**), while also attenuating the LPS-downregulated genes (Fig. 4 B, *c*, **ii**). Once again, a small group of LPS stimulated genes (<200) remains activated in statin-treated macrophages, more than cells treated with LPS alone (Fig. 4 B, *c*, **iii**), once again ruling out the possibility of overall defective LPS/TLR4 signaling. Thus, reducing cholesterol levels by statin or MCD in resting macrophages appears, on one hand, to dampen pro-inflammatory elements and, on another, to keep homeostatic function from being shut down in LPS-activated macrophages, both of which should favor later inflammation resolution.

In summary, we have identified a mechanism by which resting macrophages establish a cholesterol-dependent epigenome, thereby preparing macrophages for future immune activation. We propose a model (Fig. 4 C) in which, by reducing cholesterol in resting macrophages, there is also a cholesterol reduction in mitochondria (Fig. 4 C, *a*). This reduces mitochondrial respiration, likely through suppressed ATP synthase activity (Fig. 4 C, *b*), leading to NF-kB activation (Fig. 4 C, *c*). Activated NF-kB then goes to nucleus to perform two separate functions: 1) physically displacing nucleosomes and binding to promoter/enhancer to trigger gene expression (Fig. 4 C, *d*); and 2) upregulating histone modification enzymes and hence modifying/stablizing the epigenome (Fig. 4 C, *e*). When encountering subsequent immune stimuli, such as LPS (Fig. 4 C, *f*), these macrophages would be predicted to produce a more restrained inflammatory response with an attenuated suppression of homeostatic gene expression (Fig. 4 C, *g*).

Perhaps most importantly, we have discovered that cholesterol levels in resting macrophages directly control JMJD3 expression through NF-kB. NF-kB-reprogrammed epigenome was recently observed to persist after initial NF-kB signalling subsided, due to relatively stable histone modifications^2^. This could contribute to the so-called “epigenetic transcription memory” in macrophages^36^. By incorporating environmental cues, past or present, macrophages are prepared for future inflammatory activation in a specific fashion. Our study also places mitochondria cholesterol at the center of NF-kB activation. The pathways of NF-kB activation by MCD/statin and by LPS likely converge at mitochondria. As mitochondria acutely sense cellular physiological and pathological conditions, the mechanism we discovered here may have much broader applications for macrophage functions.

## SUPPLEMENTARY FIGURES

**Supplementary Figure 1.**
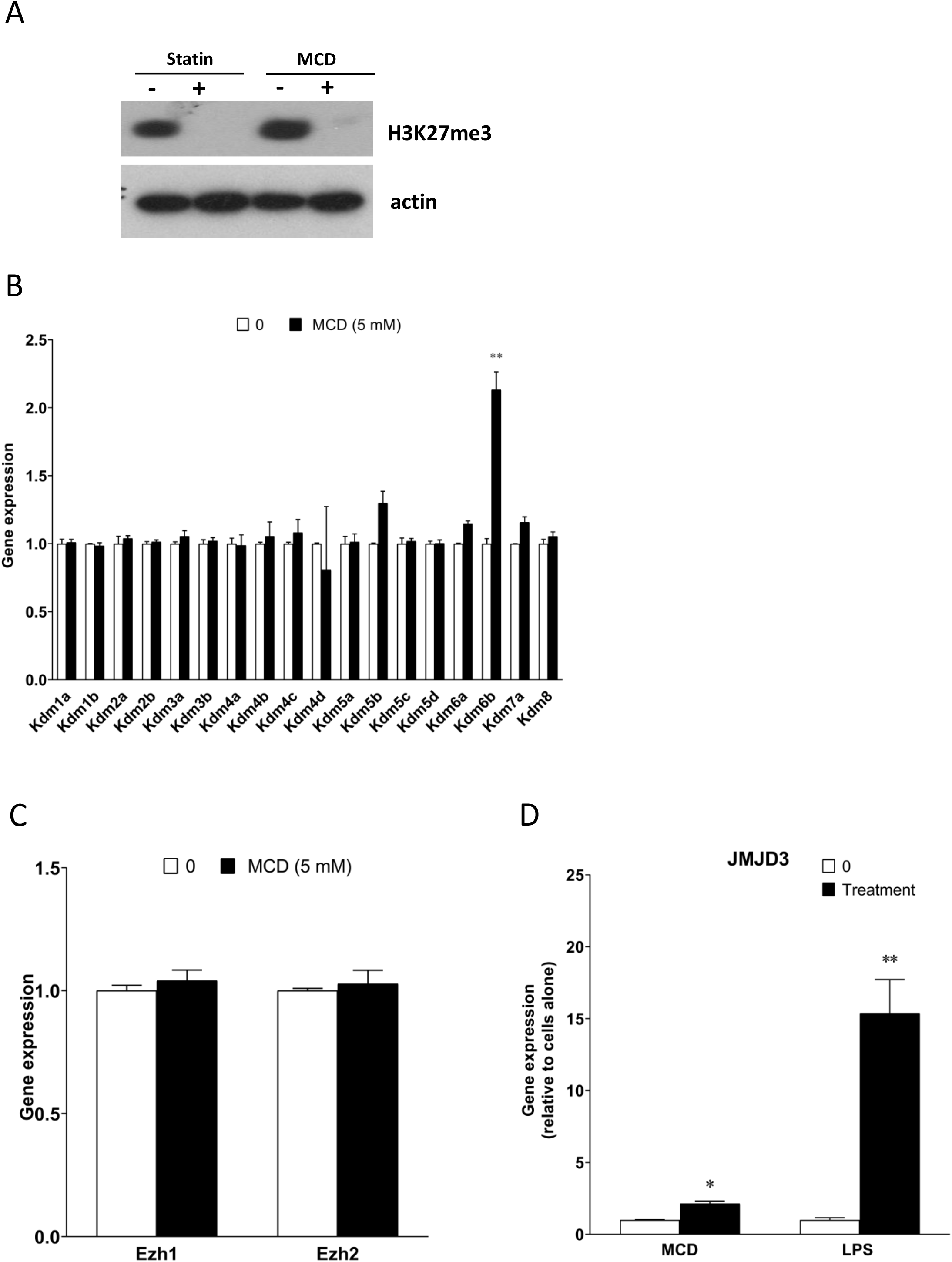
Cellular cholesterol contents regulate specific histone demethylase expression. (**A**) Western blot results of H3K27Me3 in cells treated with statins (2 d) or MCD (1h). Actin was used as a loading control. (**B**) Histone demethylases expression between MCD-treated cells and control cells. (**C**) Ezh1 and Ezh2 gene expression MCD-treated cells and control cells. (**D**) JMJD3 gene expression in MCD-, or LPS-treated macrophages. Data are presented as mean ± SD. Single asterisk (*) or double asterisks (**) indicate a significant difference with p<0.05 and p<0.001, respectively.

**Supplementary Figure 2.**
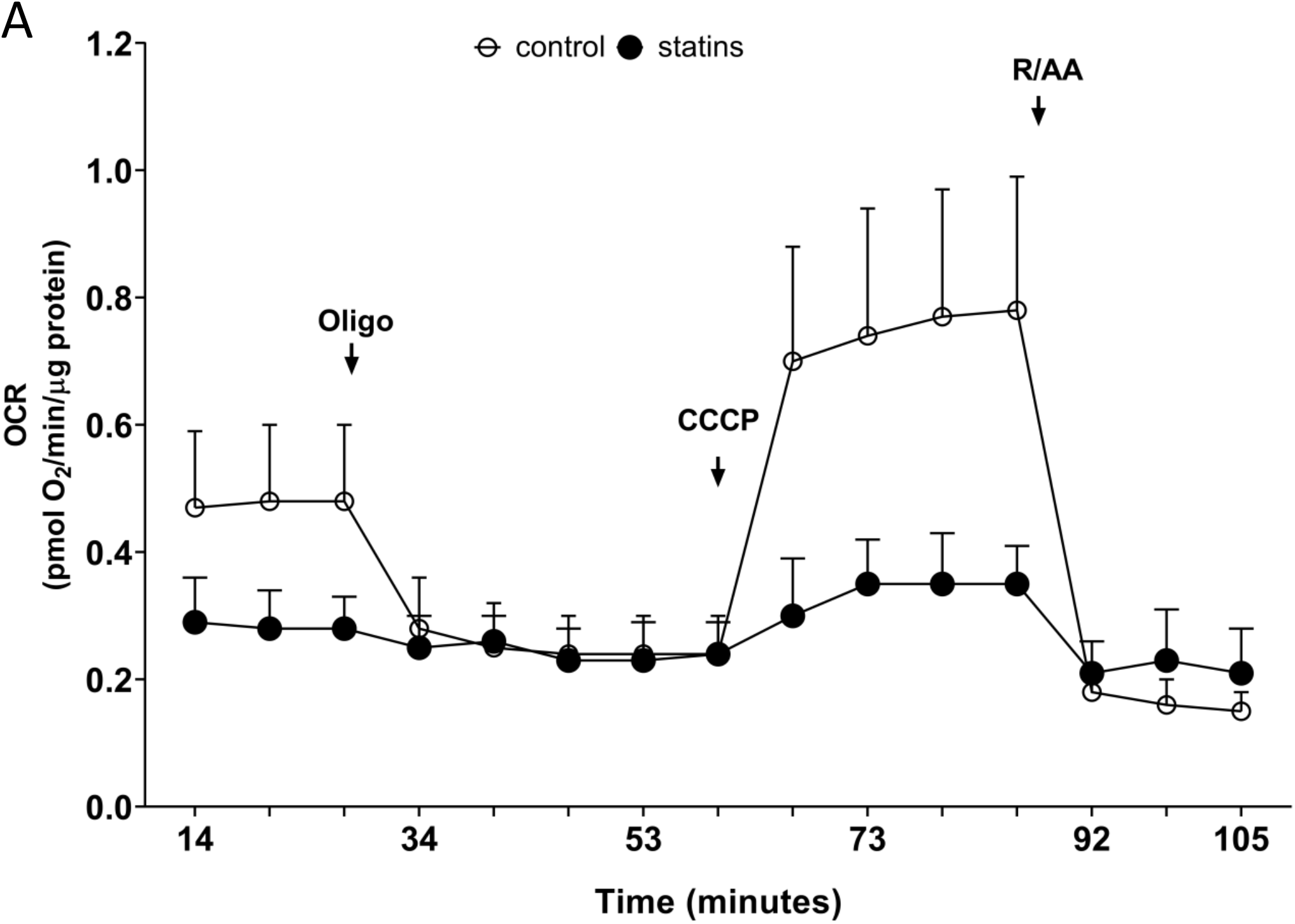
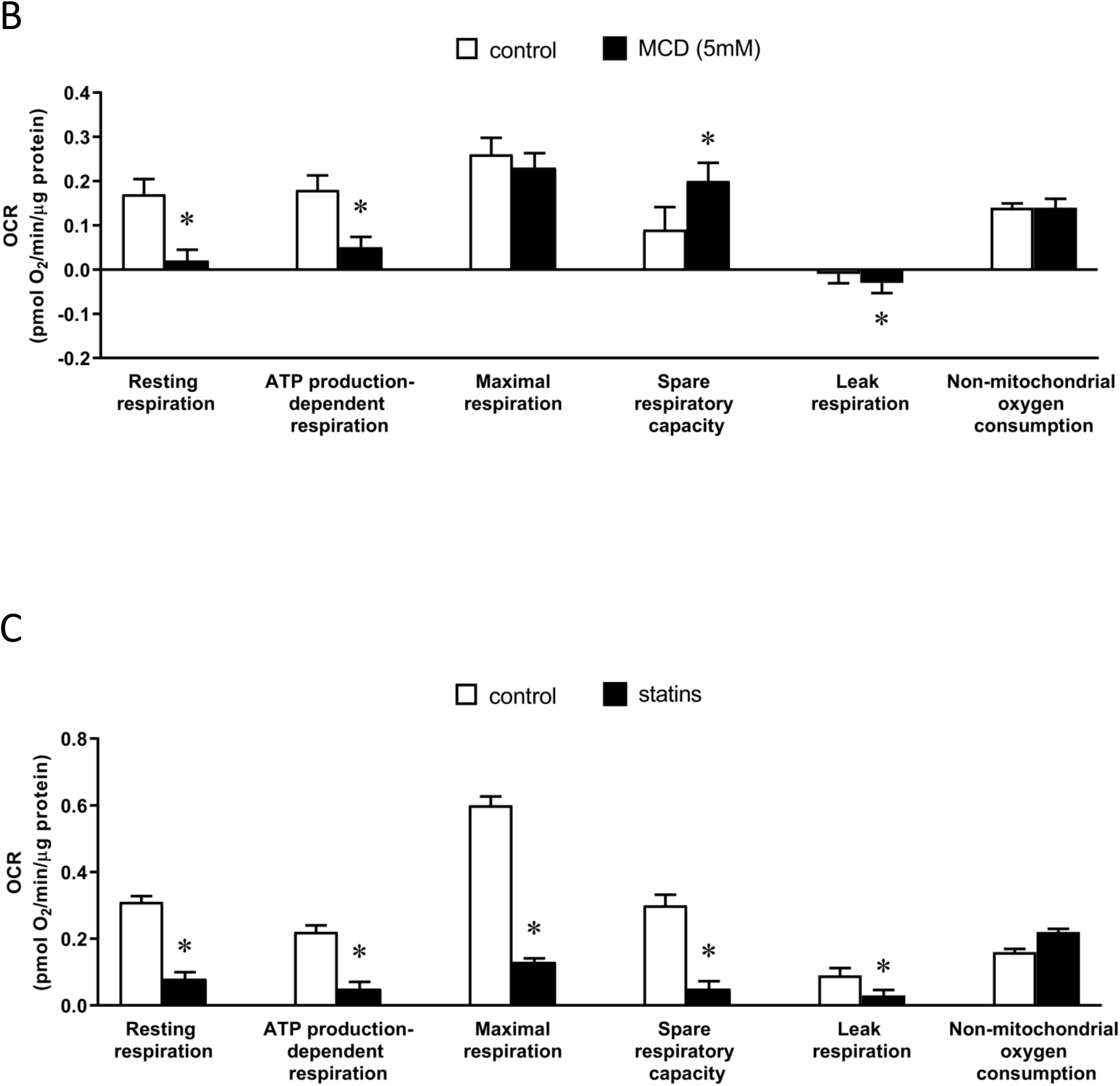
statin treatment decreases mitochondrial oxygen consumptions rates (OCR) RAW 264.7 macrophages treated with statin (2 d, 7 μM statin +200 μM mevalonate); (**A**) OCR trace of cells pretreated with or without statins; (**B**) OCR of mitochondrial and non mitochondrial parameters (functional components) in control and MCD-treated macrophages; (**C**) OCR of mitochondrial and non mitochondrial parameters (functional components) in control and statin-treated macrophages.

**Supplementary Figure 3.**
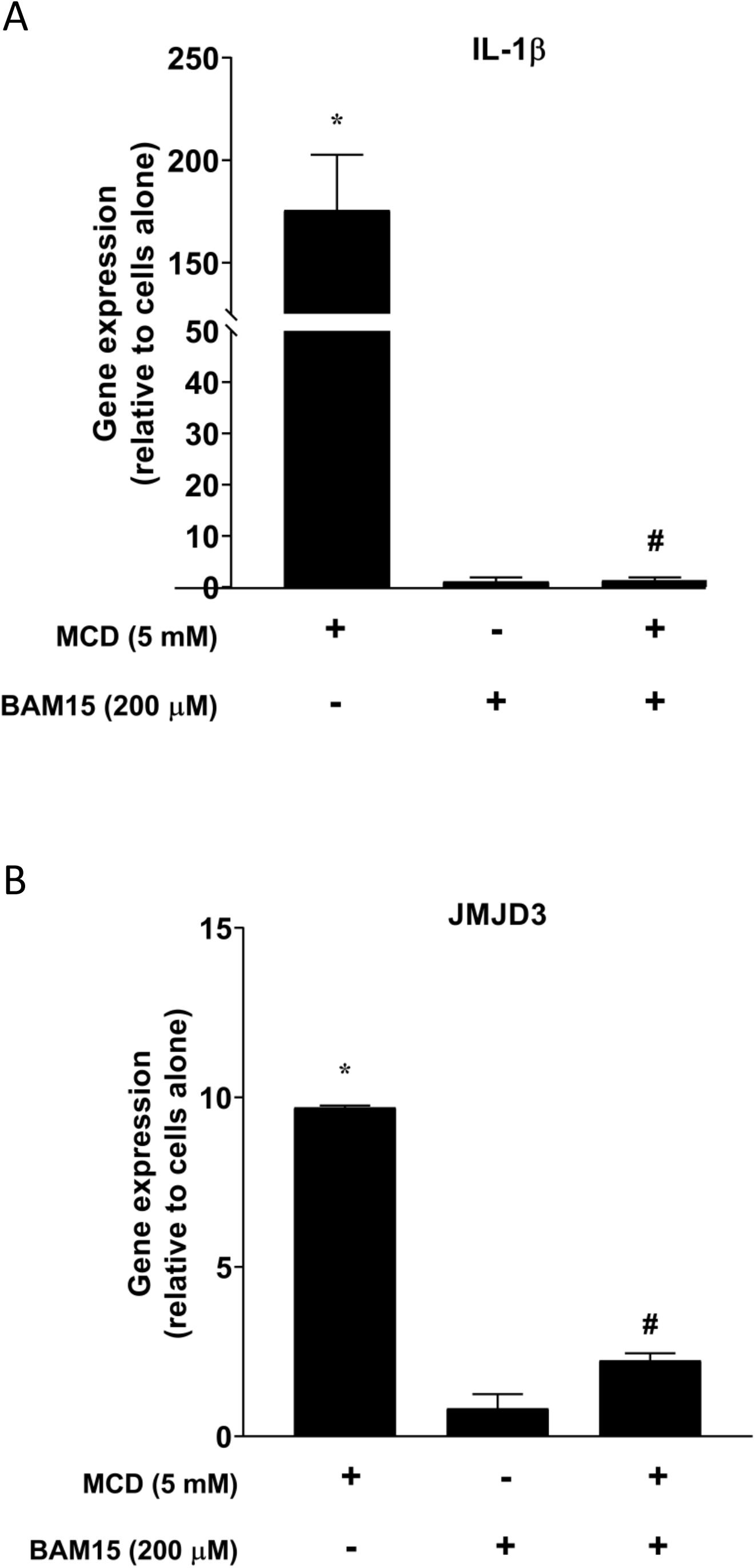
Cholesterol modulates proton flux in mitochondria to activate NF-kB pathway. Effect of BAM15 (200 μM) on Il-1β (**A**) and JMJD3 (**B**) gene expression in MCD-treated cells. Data are representative of 3 independent samples per condition and are presented as mean ± SD. Statistical analysis was performed using unpaired, two-tailed Student’s t-test. asterisk (*) and hashtag (#) indicate a significant difference with p<0.05.

## Materials and Methods

### Reagents and chemicals

Cell culture Dulbecco’s Modified Eagle Medium (DMEM) was purchased from Gibco (Thermo Fisher Scientific, Watham, MA). Antibiotics (penicillin and streptomycin) and fatty acid (FA)-free bovine serum albumin (BSA) were purchased from Sigma-Aldrich (St Louis, MO). FBS (Optima) was purchased from Atlanta Biologicals (R&D systems; Minneapolis, MN). As for the chemicals, the following inhibitors: MG-132 (proteasome inhibitor), BAY11-7082 (IKK inhibitor), and BAM15 (another mitochondrial uncoupler) were purchased from TOCRIS chemicals (part of R&D systems). Methyl-β-cyclodextrin (MCD), simvastatin, mevalonate, oligomycin, rotenone, antimycin A, carbonyl cyanide 3-chlorophenylhydrazone (CCCP), 4-(2-hydroxyethyl)-1-piperazineethanesulfonic acid (HEPES), and phosphate-buffered saline (PBS) were purchased from Sigma-Aldrich. For the IMVs study, magnesium chloride (MgCl_2_), 2-(N-morpholino) ethanesulfonic (MES), sucrose, hexane, bovine serum albumin (BSA), sulfuric acid (H_2_SO_4_), hydrochloric acid 37% (HCl), sodium chloride (NaCl), potassium chloride (KCl), dibasic potassium phosphate (K_2_HPO_4_), sodium hydroxide (NaOH), Hydrogen peroxide (H_2_O_2_), trichloroacetic acid (TCA), adenosine 5’-triphosphate disodium salt hydrate (Na_2_ATP), adenosine 5’-diphosphate sodium salt (Na_2_ADP), and cholesterol were also supplied by Sigma-Aldrich. Ammonium molybdate (VI) tetrahydrate and L-ascorbic acid were purchased from Acros Organics (part of Sigma Aldrich). Ultrapure water was produced from a Milli-Q unit (Millipore, conductivity lower than 18 MΩ cm). Rabbit anti-mouse JMJD3 primary antibody was lab-generated as described in^1^. The following antibodies were acquired from several vendors: mouse monoclonal anti-β-actin antibody (A1978) from Sigma-Aldrich, rabbit anti-mouse pCREB (87G3), rabbit anti-mouse CREB (86B10), and rabbit anti-mouse Tri-Methyl-Histone H3 (Lys27) antibodies were from Cell Signaling technology (Danvers, MA). The secondary antibody HRP-conjugated anti-rabbit antibody was from Cayman chemicals (Ann Arbor, MI). The Enhanced chemiluminescence (ECL) solutions for the Western blotting system were from GE Healthcare (Chicago, IL). The protease and phosphatase inhibitor cocktails were purchased from Sigma-Aldrich.

### Cells

The RAW 264.7 murine macrophage cell line (American Type Collection Culture [ATCC]; Manasas, VA) was maintained in 100-mm diameter tissue culture-treated polystyrene dishes (Fisher Scientific, Hampton, NH) at 37°C in a humidified atmosphere of 95% air and 5% CO_2_. The cells were cultured in DMEM-based growth medium containing glutamine and without pyruvate, supplemented with 10% [v/v] heat-inactivated fetal bovine serum (FBS) and 1x Penicillin/ streptomycin). For experiments (and routine subculture), cells were collected in growth media after 25 minutes incubation with Accutase (Sigma Aldrich, St Louis, MO) at 37°C in a humidified atmosphere of 95% air and 5% CO_2_. In preparation for experiments, cells were seeded in 6-well tissue culture-treated polystyrene plates (Fisher Scientific, Hampton, NH) at 30,000 cells/cm^2^ for 2 days until cells have recovered in time for treatments. Unless otherwise indicated, cell treatments were prepared in DMEM supplemented with 0.25% (w/v) FA-free BSA, that was sterilized by filtration through 0.2-μm pore size cellulose acetate syringe filters. For treatments, cells were washed with pre-warmed sterile PBS and were treated with media containing MCD (5 mM), cell culture-grade water (negative control). Then, the cells were incubated for 1 hour under cell culture conditions. RAW 264.7 macrophages were pre-incubated for 5 minutes with CCCP (50 μM), 30 minutes with BAM15 (200 μM), or 1 hour with NF-kB inhibitors (MG-132 [5 μM] or BAY11-7082 [10 μM]).

### RNA purification and cDNA synthesis

After treatment, the cells were lysed and collected in TRIzol (ThermoFisher Scientific, Watham, MA), and frozen at −80°C. Total RNA was extracted by phenol–chloroform extraction, followed by ethanol precipitation. RNA was purified through columns supplied in the Molecular Biology kit (BioBasic Inc., Markham, ON), according to the manufacturer’s instructions. The RNA concentrations and purity were determined using a NanoDrop One (ThermoFisher Scientific, Watham, MA) spectrophotometer. cDNA synthesis was performed on the Bio-Rad T100 PCR Gradient Thermal Cycler using the QuantiTect® Reverse Transcription kit (Qiagen, Germantown, MD), following manufacturer’s instructions.

### Reverse transcriptase quantitative PCR (RT-qPCR)

Gene expression was analyzed by real time reverse transcriptase quantitative PCR (RT-qPCR) according to the Fast SYBR Green protocol with the AriaMx real-time PCR detection system (Agilent technologies, Santa Clara, CA). Primers were ordered from Invitrogen (ThermoFisher Scientific, Watham, MA) and are listed in Table 1. Each condition was prepared in triplicates and each sample was loaded as technical triplicates for each gene (target or reference) analyzed. The mRNA levels of mouse HPRT1 or GAPDH were used as internal controls (reference gene) as indicated.

**Table 1:**
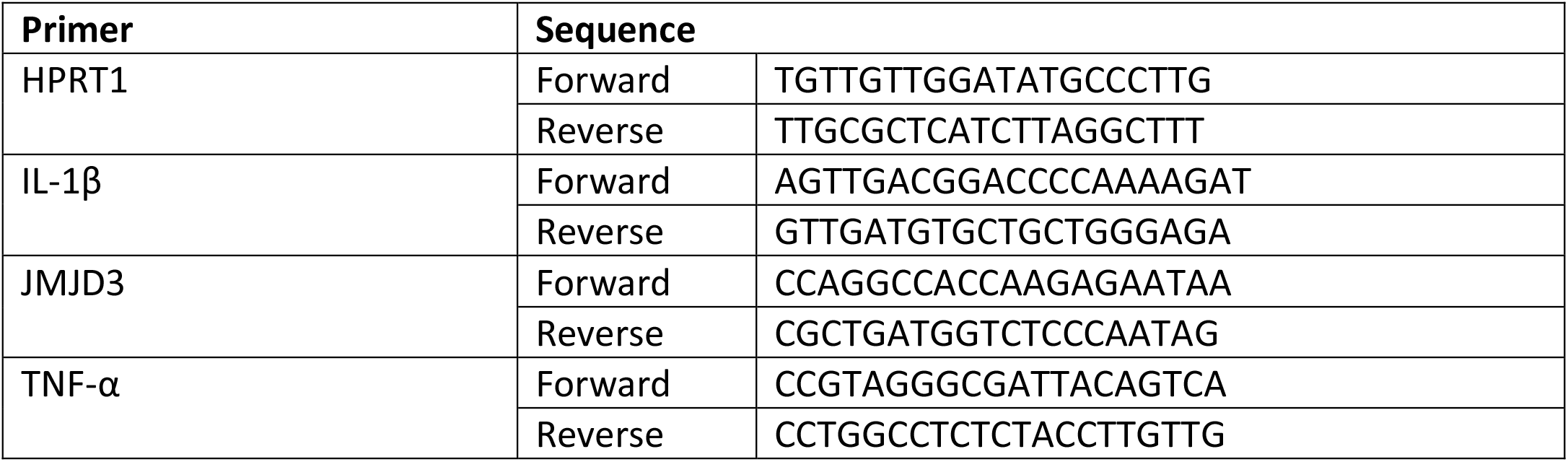
Mouse primers used for real time RT-qPCR

### Immunoblotting

For Western blot analysis, MCD-treated RAW 264.7 cells were washed in ice-cold PBS, lysed in radioimmune precipitation assay (RIPA) buffer (150 mM NaCl, 1% Nonidet P-40, 1% sodium deoxycholate, and 25 mM Tris (pH 7.6)) supplemented with a cocktail of protease and phosphatase inhibitors. Total cell lysis was achieved through sonication for 20 seconds at 20% amplitude and samples were stored at −80°C. Protein concentration was determined using the Bradford assay (Bio-Rad, 5000006). SDS buffer (313 mM Tris (pH 6.8), 10% SDS, 0.05% bromophenol blue, 50% glycerol, and 0.1 M DTT) was added, and the samples were boiled at 100 °C for 5 min. Proteins were separated by SDS-PAGE on 10% acrylamide gels. After separation, proteins were transferred onto PVDF membranes and were blocked in 5% milk powder (in PBS, 1% Triton X-100) for 1 h. Membranes were incubated with the primary antibodies in the following conditions: overnight incubation with the anti-JMJD3 antibody (1:400) at 4°C, and 1h with the anti-H3K27Me3 antibody at room temperature. After washes, blots were further incubated with an HRP-conjugated anti-rabbit antibody (1:10 000) for 1h. Blots were imaged using ECL-based film detection system.

### Cellular Oxygen Consumption

Cellular oxygen consumption rates (OCR) were measured using an extracellular flux analyzer (Seahorse XF96e; Agilent Technologies, Santa Clara, CA). Briefly, the cartridge sensors were incubated overnight at 37 °C in a hydration/ calibration solution (XF Calibrant; Agilent Technologies). Specially designed polystyrene tissue culture-treated 96-well microplates with a clear flat bottom (Seahorse XF96 V3 PS Cell Culture Microplates; Agilent Technologies) were then seeded with 80 μl of cell suspension (4.4×10^5^ cells/ml) per well and incubated 2–3 h under cell culture conditions to allow cell attachment. For treatment with MCD, cells were washed with pre-warmed sterile PBS and the culture supernatants were replaced with medium containing MCD (5 mM). Then, the cells were incubated for 1 h under cell culture conditions. For treatment with statins, cells were washed with pre-warmed sterile PBS and the culture supernatants were replaced with medium supplemented with lipoprotein-deficient serum (LPDS), containing statins (7 μM lovastatin + 200 μM mevalonate). Then, the cells were incubated for 2 days under cell culture conditions.

At the end of either treatment, the cells are washed and incubated 45 min at 37 °C in extracellular flux analysis medium (sodium bicarbonate-, glucose-, phenol red-, pyruvate-, and glutamine-free modified DMEM [Sigma-Aldrich, catalog no. D5030] freshly supplemented with cell culture-grade D-glucose (4.5 g/L), cell culture-grade L-glutamine [4 mM; Wisent], and 4-(2-hydroxyethyl)-1-piperazineethanesulfonic acid [HEPES; 4.5 mM; Sigma-Aldrich], pH adjusted to 7.35-7.40 at room temperature). OCR were measured to assess resting respiration, followed by ATP production-dependent respiration, maximal respiration, and non-mitochondrial oxygen consumption, after sequential injections of oligomycin (an ATP synthase inhibitor) at 1μM (final concentration), CCCP (an ionophore acting as a proton uncoupler) at 2μM, and rotenone together with antimycin A (complex I and complex III inhibitors, respectively), each at 0.5μM. mitochondrial ATP production-dependent respiration was calculated by subtracting the lowest OCR after oligomycin injection from resting OCR. For each parameter, OCR measurements were performed at least three times at 6-min intervals. Each condition was performed in 7–8 replicates samples. All OCR measurements were corrected for the OCR of cell-free wells containing only medium. Upon completion of the OCR measurements, the cells were washed once with PBS and lysed in 1M NaOH (40 μl/well). The lysates were kept at 4 °C for up to 24 h, and protein determination was performed using the Bradford colorimetric assay with BSA as the standard protein (Thermo Scientific). Absorbance was measured at a wavelength of 595nm using a hybrid microplate reader (Biotek).

### 27-Hydroxycholesterol (27-HC) quantification

Cell pellets were collected from RAW264.7 macrophages treated with 1, 3 and 5 mM of MCD for 1 hour. Samples were sent to the University of Texas (UT) Southwestern Medical center for analysis of 27-hydroxycholesterol based on methods previously described^2^. Briefly, lipids were extracted from macrophage samples using a modified Bligh and Dyer extraction procedure with a 1:1:1 mixture of methanol, dichloromethane, and PBS. A pool of free oxysterols was generated using base hydrolysis. Dried extracts were subjected to solid phase extraction to further isolate oxysterols. The dried lipid extracts were reconstituted in 90% methanol. Levels of oxysterols levels were measured with isotope dilution mass spectrometry using a mixture of primary and deuterated standards (Avanti Polar Lipids) added to the sample prior to extraction as a standard for quantification. Lipid extracts were resolved and detected using high performance liquid chromatography (HPLC) coupled to a triple quadrupole mass spectrometer (MS) through an electrospray ionization interface.

### Purification and formation of *E. coli* inner membrane vesicles (IMVs)

*E. coli* inner membranes were purified from *E. coli* MG1655 strain by sucrose gradient ultracentrifugation (30 min at 80,000 rpm, Optima™ MAX-XP) as previously described^3^. The *E*.*coli* inner membranes were then resuspended in either the synthesis buffer (MES 100 mM pH 6 and NaCl 25 mM) or the hydrolysis buffer (HEPES 25 mM pH 8) and ultrasonicated for 15 min (5 seconds on-off cycles at 30% power, VCX500 Ultrasonic Processors). The resulting homogeneous solution contained the inner membrane vesicles (IMVs).

### Loading and depletion of cholesterol in IMVs

Prior to any treatment, the protein concentration of IMVs was determined by the BCA protein assay and IMVs were diluted to a final protein concentration of 0.5 mg/ml. IMVs were then loaded with cholesterol using a modified protocol based on the MCD/cholesterol complex^4^. To load IMVs, 25 mM MCD were mixed with 2.5 mg/ml of cholesterol on HEPES (25 mM, pH7.8) following the protocol described in^5^. After incubation with MCD-cholesterol complexes, IMVs were ultracentrifuged (80,000 rpm, 30 min) and the supernatant removed. The IMVs were then resuspended in HEPES 25 mM (pH 7.8). After loading, the lipid, cholesterol and protein concentration of IMVs were determined by the phosphate determination assay^6^, Amplex Red cholesterol assay kit^7^ and BCA protein assay^8^, respectively. To achieve vesicles with different cholesterol content, cholesterol-loaded IMVs (1mg/mL protein concentration) were treated with increasing MCD concentrations (from 0 to 7.0 mM) for 30 min at 37^o4,9^. Cholesterol-depleted vesicles were ultracentrifuged (80000 rpm, 30 min) and the pellet was resuspended in HEPES (25 mM, pH7.8). Lipid, protein and cholesterol concentration of depleted samples were quantified again after the MCD treatment.

### Hydrolysis and synthesis of ATP on cholesterol-doped IMVs

ATP hydrolysis was performed by adding a total concentration of 2 mM ATP to 200 mM IMVs (lipid concentration) and incubated for 30 minutes. The concentration of phosphates from ATP hydrolysis was measured using the malaquite green assay^10^. ATP synthesis was triggered by promoting a ΔpH across the IMV membranes by mixing the samples (IMVs resuspended in MES 100 mM pH 6) with an external buffer with higher pH and in the presence of inorganic phosphate, ADP and magnesium ion (HEPES 100 mM pH 8, 5 mM Pi, 2.5 mM ADP, 5 mM MgCl2, 25 mM NaCl) at a volume ratio of 1:10 and incubated for 2 to 5 minutes. The reaction was stopped by adding 20% TCA at a ratio volume of 10:1, and then the samples was equilibrated to neutral pH^11^. ATP concentration after synthesis was measured using ATP detection assay kit (Molecular Probes) with a luminometer GloMax®-Multi Detection. Amplex™ Red Cholesterol Assay Kit and Pierce® BCA Protein assay kits were supplied by ThermoFisher. Luminescent ATP Detection Assay Kit (based on firefly’s luciferase / luciferin) was purchased from Molecular Probes.

### RNA-sequencing

RAW 264.7 macrophages in duplicates were either left untreated or treated with 5 mM MCD for 1 hour. Total RNA was purified as described above, and the RNA concentrations and purity were determined using a NanoDrop One (ThermoFisher Scientific, Watham, MA) spectrophotometer. Library preparation and 150-bp paired-end RNA-Seq were performed using standard Illumina procedures for the NextSeq 500 platform. The RNA-Seq data were analyzed using the Bowtie, TopHat, CuffLinks, and CummeRbund software suite^12^. Briefly, reads were mapped to the mm9 genome (University of California, Santa Cruz, California, USA) and known mm10 transcripts (RefSeq mm10 build37.2) using Bowtie version 2.2.4^13^ with default parameters. Splice junctions were determined from the mapped reads using TopHat version 2.0.12^14^ with default values. Transcript quantification was performed with HT-Seq version 0.6.1^15^. Identification of differentially expressed genes was performed with the edgeR Bioconductor package^16^. RNA-Seq data were deposited in the UCSC genome browser database.

### ATAC-seq

RAW 264.7 macrophages in duplicates were either left untreated or treated with 5 mM MCD for 1 hour. Cells were detached using 0.5% trypsin and resuspended in chilled 1X PBS containing 1 mM EDTA. Visible cell numbers per sample were obtained by staining the cells with trypan blue and counting them using hemocytometer. 50,000 cells were used for performing ATAC-Seq. Cells were washed in 100 μl of ice cold 1X PBS and centrifuged at 500 x g for 5 minutes. Nuclei are prepared by lysing the cells in ice cold lysis buffer containing 10 mM Tris-Cl, pH7.4, 10 mM NaCl, 3 mM MgCl2 and 0.1% IGEPAL CA-630. Nuclei were pelleted by spinning the samples at 500 x g for 10 minutes in fixed-angle cold centrifuge and proceeded for tagmentation reaction. A 25 μl tagmentation reaction was setup by resuspending the nuclei in 12.5 μl of 2x Tagment DNA buffer and 5 μl of TDE1 transposase. Samples were incubated at 37°C for 30 minutes with gentle intermittent mixing. Following transposition, the sample volume was made up to 50 μl using resuspension buffer and were processed for DNA preparation. For tagmented DNA clean-up, 180 μl of Zymo DNA binding buffer was added and mixed thoroughly before loading on to Zymo-spin concentrator-5 columns. For transposase free DNA, the samples were eluted in 16.5 μl of elution buffer. Purified tagmented DNA was amplified with KAPA HIFI polymerase (12 PCR cycles) and Unique Dual Primers from Illumina (Cat number 20332088). Size-selection (L:1.1; R :0.6) was performed with KAPA Pure beads and size distribution of the final libraries was assessed on bioanalyzer (Agilent). Libraries were quantified by qPCR and loaded equimolarly on a S4 Novaseq flowcell. Each library was sequenced with a coverage of 50M paired-end reads (PE100).

### ATAC-seq analysis

FastQC^17^ was used to perform preliminary quality-control for paired-end FASTQ reads retrieved for control and MCD-treated samples (two replicates each) respectively. Sequencing adapters were trimmed from the reads using Cutadapt v2.6^18^ and aligned to the mouse genome (GRCm38-mm10) using Bowtie2 v2.4.4^13^. The resulting alignments were filtered using Samtools v1.10^19^ to retain those that were mapped in proper pairs (-f 2) and based on their MAPQ scores (-q 10). Finally, duplicates were removed using Picard (java v1.8.0) and the resulting filtered alignments were converted to genome-browser compatible 6igwig files with CPM normalization using bamCoverage (deeptools v3.5.1)^20^. The filtered alignments were concurrently also used to identify (narrow) peaks for each of the replicates using MACS3 v3.0.0^21^ with default parameters. Differential peaks between control and MCD-treated samples were identified using DiffBind^22^ and functional annotation of peaks was performed using ChIPseeker^23^.

### Statistical analysis

Statistical analyses between data groups were performed with PRISM software (GraphPad). Data for real time RT-qPCR and Seahorse experiments are presented as the mean ± S.D. as indicated. The statistical significance of differences between groups was analyzed by Student’s *t* test. Differences were considered significant at a *p* value < 0.05.

## Footnotes

* This particular assay could detect relative changes, but not absolute amount of cholesterol in IMM24.

